# Near-infrared spectroscopy-based models correctly classify *Abies alba* seed origin and predict germination properties

**DOI:** 10.1101/2025.06.09.658572

**Authors:** Jeanne Poughon, Camille Lepoittevin, Eduardo Vicente, Marion Carme, Georgeta Mihai, Francisco Lario Leza, Andrea Piotti, Camilla Avanzi, Maurizio Marchi, Giovanni Giuseppe Vendramin, Caroline Scotti-Saintaigne, Bruno Fady, Caroline Teyssier, Marta Benito Garzon

## Abstract

Forestry industry requires high-quantity and quality seeds for afforestation and assisted migration programs. Finding reliable non-destructive methods to characterize seeds would significantly enhance efforts to identify climate-adapted populations. This study presents near-infrared (NIR) spectroscopy models to classify seed origin and predict germination characteristics at different temperatures non-destructively. We focus on *Abies alba* Mill., a key European forest tree with genetic variation along climatic gradients and seeds with shallow physiological dormancy. Seeds from six populations were analyzed using NIR spectroscopy, and germination was tested at 15°C, 20°C, and 25°C after stratification treatments at 4°C (0 or 3 weeks). Population classification accuracy using Partial Least Squares Discriminant Analysis was 69%, with significant NIR peaks at 1712, 1929, and 2111 nm, linked to moisture content and storage compounds. NIR spectra explained 51% and 65% of the variation in germination probability and timing using Partial Least Squares Regression, with significant peaks at 1712, 1929, 2111, 1632, and 2073 nm. General Linear Mixed-Effects Models showed that a NIR predictor contributed to 39% of the germination probability variance explained by fixed-effects, and the stratification treatment was the most important driver explaining germination time. Our results proved the utility of NIR-based tools to effectively classify bulked seeds and predict germination, opening new perspectives to nursery and forestry sectors and populations’ adaptation and adjustments to warming climate. This study will facilitate further investigations on the physiological processes that occur during dormancy, a critical process for forest regeneration given the expected impact of shorter and warmer winters on seed behavior.

## 1. Introduction

Phenotypic plasticity and local adaptation are the two main evolutionary processes maintaining fitness variation across species ranges. These two processes are relatively well studied across species ranges in adult forest trees. Conversely, much less is known for young and early stage trees (Leites and Benito Garzón, 2023), and particularly for seed traits such as germination-related ones (Vicente and Benito Garzón, 2024). Yet, differences in tree seed composition and characteristics in terms of germination and dormancy can be found across provenances, mother plants and years of collection (Alonso-Crespo et al., 2020; Lacerda et al., 2004; Loewe et al., 2017; Soriano et al., 2011; Vaughn et al., 2022). This knowledge gap is crucial when assessing the impact of climate change on early developmental stages. Tree seed germination traits, such as germination probability and germination time, combine a significant impact on individuals’ fitness (Verdú and Traveset, 2005), and a pronounced climatic sensitivity (Walck et al., 2011). A decrease in germination rate could constitute a major bottleneck for recruitment (Tiebel et al., 2023), limiting forest regeneration and establishment, thereby slowing range shifts (Kroiss and Hille Ris Lambers, 2015).

Germination is strongly tied to seed composition and especially water content and storage compounds (Nonogaki et al., 2010), which vary with environmental conditions and will be affected by climate change (Durr et al., 2018; Fenner, 2010; Moreira et al., 2024; Pesendorfer et al., 2020). Germination’s response to climate change is even more uncertain when considering winter warming for species that require specific cold conditions beforehand to release their physiological dormancy and germinate (Walck et al., 2011; Willis et al., 2014).

The study of seed trait variation, particularly dormancy and germination, is complex, time consuming, and costly. Associating seed composition characteristics to dormancy and germination traits is challenging because biochemical analyses are destructive, whereas germination and dormancy experiments require the monitoring of seeds over long periods. Additionally, biochemical analyses demand large sample sizes, which are often not available when studying natural populations with masting years (i.e., seed production only occurs every two to four years, as in the case of silver fir). Hence, understanding seed germination in interaction with dormancy and seed composition would highly benefit from non-destructive, cost-effective techniques. Furthermore, an increasing demand for Forest Reproductive Material (FRM) of high quality and quantity for afforestation by forestry and nursery industries is expected (Mataruga et al., 2023), on par with the European Union (EU) Forest Strategy for 2030 that aims to plant three billion additional trees. To meet this objective in a context of climate change, the European forest sector needs to make an appropriate use of Forest Genetic Resources (FGR), as planting climate-adapted populations can increase the probability of trees to survive as well as increase carbon sink (Chakraborty et al., 2024). Therefore, finding a reliable method to obtain relevant information from stored seeds — such as origin, germinability, germination phenology, or dormancy status —is crucial, not only to bridge existing knowledge gaps, but also to meet future demands of afforestation goals.

Near-infrared reflectance (NIR) spectroscopy is a low-cost, non-destructive, fast and reliable technique to study characteristics linked to chemical composition and physical properties of biological material. The near-infrared spectrum lies between the visible and mid-infrared regions 100 (780nm to 2526nm). When NIR radiations interact with an organic sample, the vibration of molecular bonds creates absorption bands that allow chemical compounds to be distinguished (Wang et al., 2022). NIR spectroscopy has previously been used to discriminate provenances in tree species, thanks to variation in storage compounds (Tigabu, 2003; Farhadi et al., 2017). Several studies have also been able to evaluate components of seed viability, including germination ability in tree species (Dumont et al., 2015; Tigabu, 2003). However, there are very few studies which used natural occurring variation in germination traits, instead of data gathered from artificially aged seeds (Çelikta and Konuşkan, 2020).

*Abies alba* Mill. (Silver fir) provides a compelling case study for exploring seed trait variations. Its seeds present a shallow physiological dormancy that requires up to six weeks of wet-cold conditions (Wolf, 2003). This evergreen conifer is distributed from low to high altitudes in the mountainous regions of eastern, western, southern and central Europe (Wolf, 2003; Mauri et al., 2016). Beyond its economic importance in forestry (Wolf, 2003), silver fir acts as a keystone species, maintaining high biodiversity in European forest ecosystems (Simberloff, 1998) and providing important ecosystem services (Ruosch et al., 2016). Silver fir shows intraspecific variation in many fitness-related traits, including those at the seed level: morphological traits (such as seed mass and size), germination probability and germination time were found to vary among provenances, both at the local scale (Boncaldo et al., 2010; Messeri et al., 1963; Mihai and Alexandru, 2021; Morar et al., 2023) and across the species range (Alt, 2022). To our knowledge, no study has used NIR to characterize provenance and germination in *A. alba* seed before.

Our main goal is to test whether the intraspecific variation in seed composition and seed germination traits can be accurately predicted by seed NIR spectra, as well as investigate the variation in the effects of dormancy release on these germination traits. More precisely we aim to: *i)* identify *Abies alba* populations using their seed NIR spectra; *ii)* investigate the potential of NIR spectra to explain population germination traits; *iii)* evaluate the effect of dormancy release on germination traits across range-margin populations that are likely adapted to local conditions. To these aims, seeds of *A. alba* were collected from six distinct provenances in autumn 2022. Among them, 4119 seeds were germinated at three different temperatures with and without stratification treatments and monitored over time and NIR spectra was measured on 2298 other seeds. We performed Partial Least Squares Discriminant Analysis (PLS-DA) on the NIRs data to discriminate between populations and applied Partial Least Square Regression (PLSR) to establish the relationship of NIR spectra with germination traits. Lastly, we analyzed the contribution of NIR spectra to explain the germination together with dormancy release (i.e., stratification) and temperature treatments (General Linear Mixed-Effects Models, GLMM).

## 2. Material and methods

### 2.1 Seed sampling

In October 2022, we collected *Abies alba* cones from six populations (Figure 1, Table 1). We selected populations at different edges of the distribution range (Caudullo et al., 2017), to maximize the possibility for different local adaptations. We also had three populations in the same mountain range (i.e., the Pyrenees) to test for local differences. Cones were dried at 20°C for one month until opening. Then, mature seeds were manually extracted, and empty seeds were sorted by airflow or manually removed. Seeds were stored at −5°C in lots by provenance and mother-tree, except for the Spanish population where there was only one bulk seed lot from 20 mother trees.

**Figure 1.**
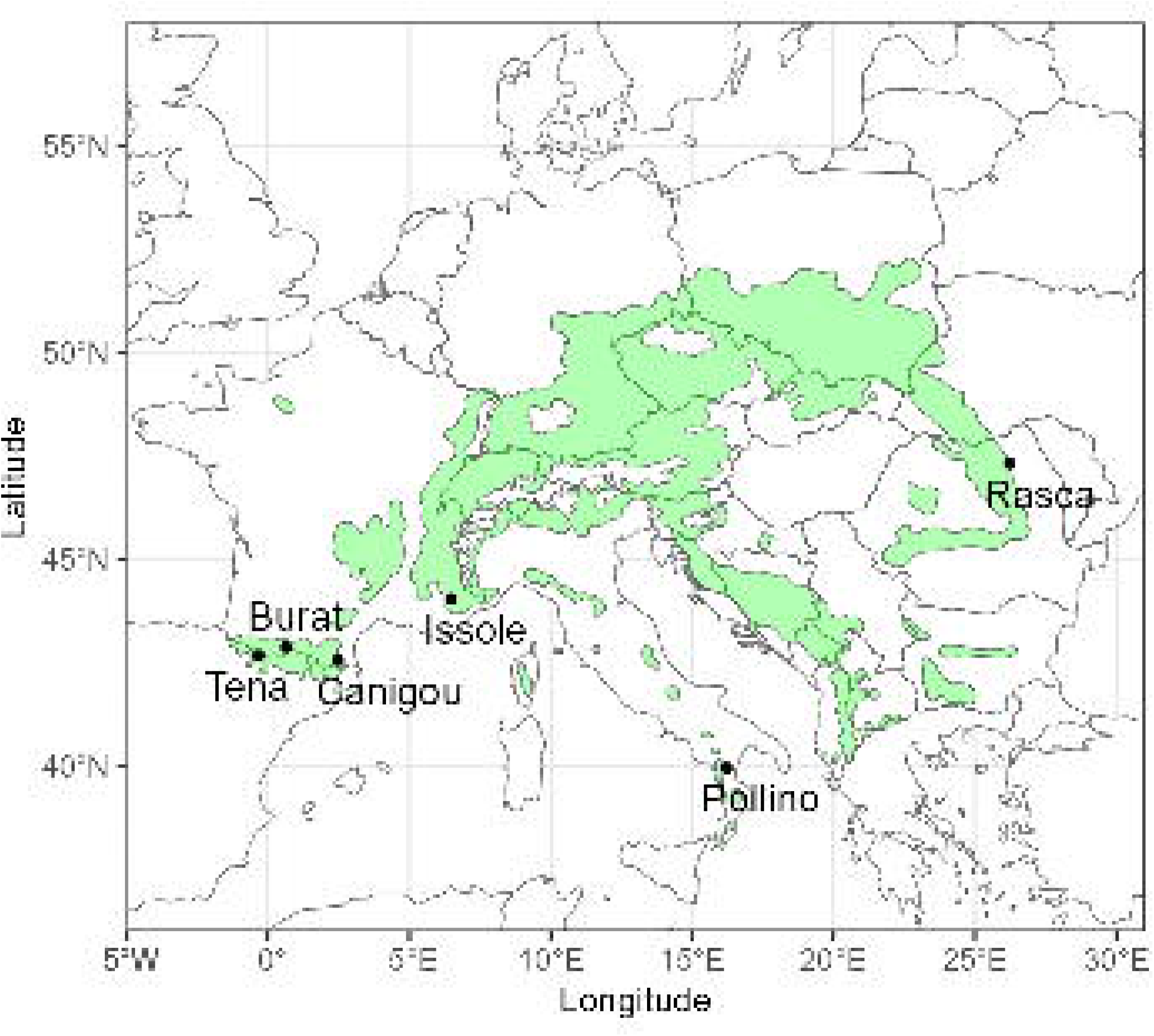
Locations of the Abies alba seeds collected for the experiment. The green range represents Abies alba distribution (Caudullo et al., 2017). The location of the population sampled is symbolized by the black points.

**Table 1.** Seed provenance. The climate data was obtained from ClimateDT (Marchi et al., 2024). Maximum spring temperature and minimal winter temperature were calculated between March and June, November and February respectively, then averaged for the period 1900-1960

| Provenance | Country | Longitude (dd) | Latitude (dd) | Elevation (m) | Maximum spring temperature (C°) | Minimum winter temperature (C°) | Temperature seasonality (standard deviation × 100) | Storage date |
| --- | --- | --- | --- | --- | --- | --- | --- | --- |
| Issole | France | 6.475 | 44.045 | 1195 | 13.26 | -5,70 | 666.76 | 11/29/2022 |
| Burat | France | 0.647 | 42.879 | 1372 | 10.68 | -4,22 | 573.92 | 12/16/2022 |
| Rasca | Romania | 25.08 | 47.2 | 1135 | 14.13 | -6.34 | 872.78 | 02/09/2023 |
| Canigou | France | 2.474 | 42.558 | 1501 | 10.82 | -4.91 | 566.25 | 12/16/2022 |
| Pollino | Italy | 16.226 | 39.951 | 1493 | 11.34 | -2.70 | 607.617 | 11/29/2022 |
| Tena | Spain | -0.336 | 42.669 | 1867 | 8.28 | -6,64 | 622.88 | 01/05/2023 |

Fresh and dry seed mass was estimated for each mother-tree as the mean mass of 20 seeds. Seeds were dried at 60°C several days until their weight did not change compared with that of the previous day to obtain the mean moisture content.

We first used the seeds in a germination experiment where we monitored germination proportion and timing. Subsequently, we measured the NIR spectra of seeds from the same seed lots used in the germination experiment whenever possible (Issole, Rasca, and half of the mother trees from Pollino); otherwise, we used seeds from the same population.

Significant differences in moisture content and dry seed mass between populations were tested with an ANOVA and a Tukey test for pairwise comparison (Supplementary Figure S1).

### 2.2 Germination experiment

To measure the variation in germination characteristics in the seeds and the combined effects of dormancy release and temperature, we performed a germination experiment in 2023 with a total of 4119 seeds. Half of the seeds were stratified for three weeks in humidified blotting paper at 4°C in a refrigerator. We opted for a stratification period of three weeks instead of the recommended six weeks (Wolf, 2003), to ensure that the full extent of stratification needs would not be met, and thereby avoid masking differences among populations. Similar stratification durations have been used in previous studies (Afsane et al., 2012; Alt, 2022; Boncaldo et al., 2010; Messeri et al., 1963).

Both stratified and non-stratified seeds were sown in 6 × 6 cm, 8 cm depth pots with 50% peat and 50% sand substrate (ten seeds per pot). The pots were placed in three climatic chambers (Snijder LABS, micro clima-series) at 15, 20 and 25°C with 70% humidity, 100 µmol m^−2^ s^−1^ light intensity, and a photoperiod of 16/8 light/dark hours. Nighttime temperatures in each chamber were set 8°C lower than the corresponding daytime temperature. The seeds came from eight mother-trees per populations except for the Italian and the Spanish ones that counted 16 and 20 mother-trees respectively (pooled together for the Spanish population). The number of seeds per experimental condition (temperature × stratification) per mother-tree varied from three to 20 depending on the quantity of seeds available (Supplementary Table S1). We watered the seeds three times a week with distilled water to maintain optimal humidity for germination.

To measure germination proportions and germination timing, we monitored seeds germination status (detection of a radicle at least 2 mm long) and germination timing (number of days between sowing and germination) thrice a week during six months.

### 2.3. NIR spectroscopy measurements

To assess the potential of NIR spectroscopy to characterize populations and explain germination performances, the NIR spectra of 2298 seeds were measured, with 48 seeds per mother tree, and eight mother trees per population. These measurements were not conducted on the same seeds used in the germination assay, as our goal was to assess whether such spectral data could be used to extrapolate information about entire seed lots, or even across different seed lots from the same provenance and year of collection. This approach was intended to better reflect potential applications by forest managers and nurseries, who are unlikely to be able to test every individual seed.

The NIR spectroscopy measurements were conducted with a benchtop Fourier transform-NIR (FT-NIR) spectrometer (MultiPurpose Analyser-MPA, Bruker Optics, Ettlingen, Germany) equipped with an optical fiber reflectance probe. Each seed was laid in an individual well from a 48-well-plate, on top of 1 ml of 2% agarose gel. The well-plate allowed an easy identification (coordinates of the seeds on the plate) and manipulation of the seeds without mistake.

The spectra were acquired by placing manually the optical fiber in contact with each seed in each plate. The following instrumental settings were applied: 9000 to 4000 cm^−1^ (1111–2500 nm) spectral range, with an optical resolution of 8 cm^−1^, 32 scans per measure and a scanner velocity of 10 kHz (probe). Opus software (v8.5, Bruker Optics, Ettlingen, Germany) was used for instrumental control and spectra acquisition.

### 2.4 Spectra preprocessing

We removed the end of the selected spectra (2273 to 2500 nm) as this area showed high variability not associated with identifiable absorption peaks, suggesting poor data quality. To correct for undesirable effects caused by seed morphology and operator (differences in spectral path length), and external factors (such as light scattering, baseline shifts, random noises) (Barnes et al., 1989), we applied several pre-treatments (Standard Normal Variate, detrending, derivation of 1^st^ and 2^nd^ order) with only the best retained for each model.

### 2.5 Partial least square discrimination analysis to classify seeds NIR spectra by population

We performed Partial Least Square Discriminant Analysis (PLSDA) to test the differences in NIR spectra between populations. PLSDA produces a linear multivariate model that relates a matrix of observation (*X*) (the spectra) and a qualitative variable (the populations) encoded in a binary (0/1) matrix for where each column represents a different class (*Y*) (Wold et al., 2001). The model finds the best group of orthogonal linear combination of the *X* variables (called the latent variables) to predict *Y*. The analysis was computed with the *rchemo* R package (Brandolini-Bunlon et al., 2023) which uses the kernel algorithm from Dayal and MacGregor (1997).

To select the best pre-treatment and determine the optimal number of latent variables, we used k-fold cross-validation with mean classification error (12 folds repeated 20 times, balanced to maintain initial proportions of mother-trees). The selected model used the pre-treatment that gave the lowest mean classification error overall and had the lowest number of latent variables within one standard deviation of the best model’s error.

The initial dataset was divided into a training set (75%; 1,723 seeds) and a test set (25%; 575 seeds) for external validation. To represent all the provenances and avoid correlations between both sets, we constituted the test set by drawing two random seed lots (*i.e.,* seeds from two mothers) from each provenance (Supplementary Table S2).

Internal and external model performances were evaluated with standard classification measurements (Ballabio and Consonni, 2013): overall accuracy (true positive/total number of predictions), sensitivity (true positives/(true positives + false negatives), specificity (true negatives/(true negatives + false positives), and balanced accuracy (sensitivity + specificity)/2. Classification predictions were used to build a confusion matrix between model predictions and the actual prediction to showcase the patterns in population misclassification.

We defined the most relevant spectral regions to identify provenances using Variable Importance for the Projection (VIP) (Wold et al., 2001). VIP summarizes the contribution of each variable to the discrimination of the groups, for both the X matrix and the Y matrix modelling. Variables with a VIP value superior to one were considered significant (Eriksson et al., 2013).

### 2.6 Partial least square regression of population germination probability and time with NIR spectra

To investigate the potential of NIR spectra to explain population germination traits, we performed two PLSR with population germination probability and time as the matrixes. The population germination probability and the population germination time were computed as the average at the population level on the non-stratified seeds at 20°C (the average temperature condition). The same NIR spectra datasets as previously was used, and pretreatment and optimal number of latent variables were chosen similarly.

As a measure of the variance explained by the models, we used the *rchemo* package to calculate the determination coefficient R^2^ on the test set.

Finally, we performed a VIP analysis as previously described in paragraph 2.5 to identify the most relevant spectral regions for predicting germination probability and time.

### 2.7 Mixed-effects models of germination probability and time

We used generalized linear mixed-effects models (GLMMs) of germination probability and germination time. Specifically, we fitted logistic mixed-effects models with a logit link function for germination status, and negative binomial mixed-effects models for germination time (*glmmTMB* R package; Brooks et al., 2017). These models served the purpose of evaluating the relevance of population NIR spectra in explaining the variation in seed germination probability and time, as well as the effect of dormancy release on these traits. Fixed-effects were dry seed mass, NIR spectra scores, climate of origin, stratification, temperature of germination, and mother-trees were random effect. Climate of origin, dry seed mass and moisture content were chosen as co-variables to weight the explanatory potential of NIR spectroscopy against other *a priori* information on seed lots.

To represent the climate of origin of the seeds in the model, we evaluated several relevant climate variables and selected the one showing the highest correlation with germination traits (Supplementary Table S3). The variable best correlated with germination probability was temperature seasonality (standard deviation of temperature x100), and the best correlated with germination time was the minimal winter temperature (mean between November and February). Both were estimated and averaged for the period 1900-1960 (data from ClimateDT, (Marchi et al., 2024)). For the population NIR spectra variables, we performed a PCA on the Standard Normal Variate (SNV) spectra and averaged the PCA scores of the first three axes by population. We decided to use Standard Normal Variate (SNV) because it allowed the secondary axes of the PCA to retain more information than other pre-treatments. This allowed us to obtain three variables that synthesize the NIR spectra at a population level. Contribution of each wavelength to the construction of the axes is shown in Supplementary Figure S2.

The general initial models took the following form:

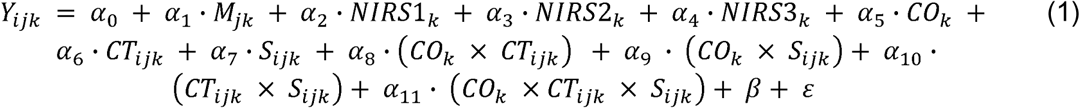

With *Y_ijk_* the trait response of the *i*th seed of the *j*th mother-tree of the *k*th provenance (either the probability to germinate or the germination time in days); *M_jk_* the mean seed mass of the *j*th mother tree of the *k*th population; *NIRS*1*_k_* the mean score of the *k*th population on the first axis of the PCA on NIR spectra; *NIRS*2*_k_* the mean score of the *k*th population on the second axis of the PCA on NIR spectra; *CO_k_* the mean climate of origin of the *k*th population in winter (November to February) averaged for the period 1900-1960; *CT_ijk_* the day temperature of the climatic chamber in which the *i*th seed of the *j*th mother-tree of the *k*th provenance was cultivated; *S_ijk_* the stratification treatment (binary) of *i*th seed of the *j*th mother-tree of the *k*th provenance; *β* the random effects and *ε* the residuals.

To select the fixed factors retained in the models we followed stepwise backward selection with a Chi square test (*drop1* function; (R Core Team, 2023)). We stopped at the most parsimonious model that still contained one NIR variable. We obtained the final following models:

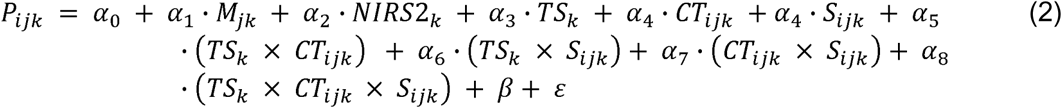

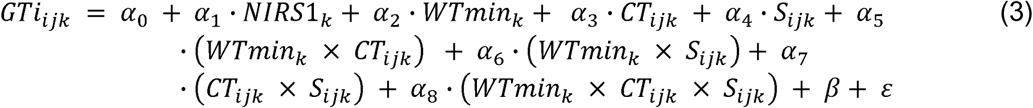

With *P_ijk_* the probability the *i*th seed of the *j*th mother-tree of the *k*th provenance will germinate; *TS_k_* the temperature seasonality of the *k*th population; *GTi_ijk_* the germination time in days of the *i*th seed of the *j*th mother-tree of the *k*th provenance; *WTmin_k_* the mean minimal temperature of the *k*th population in winter (November to February).

To analyze the variance explained by the models and by each variable selected, we computed Nakagawa’s R^2^ for binomial (theoretical R^2^) and negative binomial models (trigamma R^2^) (Nakagawa et al., 2017) with the function *r.squaredGLMM* from the *MuMin* R package (Kamil Bartoń, 2010), as well as a commonality analysis. Commonality analyses allow to decompose R^2^ and indicate how much variance is explained by a single predictor (unique effect) and how much variance a predictor shares with all possible combinations of the other predictors (common effects) (Ray-Mukherjee et al., 2014).

Moisture content was not included in the model selection presented here because of its high correlation (>0.5) with *NIRS*2. Supplementary material includes a model obtained after model selection with moisture content to discuss the interchangeability of moisture content and *NIRS*2 (Supplementary Equation S1).

## 3. Results

A summary of the mean characteristics for each population, stratification treatment and chamber temperature, is shown in Supplementary Table 4.

The PLSDA performed on seeds spectra to classify them in different populations had 35 latent variables according to cross-validation with the one standard error rule (as defined in paragraph 2.5). The classification accuracy on the test set was 69%, which is 27% lower than on the training set (Table 2). The model performed differently on the test set depending on the population, with some populations better identified than others. Issole and Tena had the highest accuracy and sensitivity, while Pollino had the lowest values for these two indicators of model performance. Issole also had the lowest specificity.

**Table 2.**
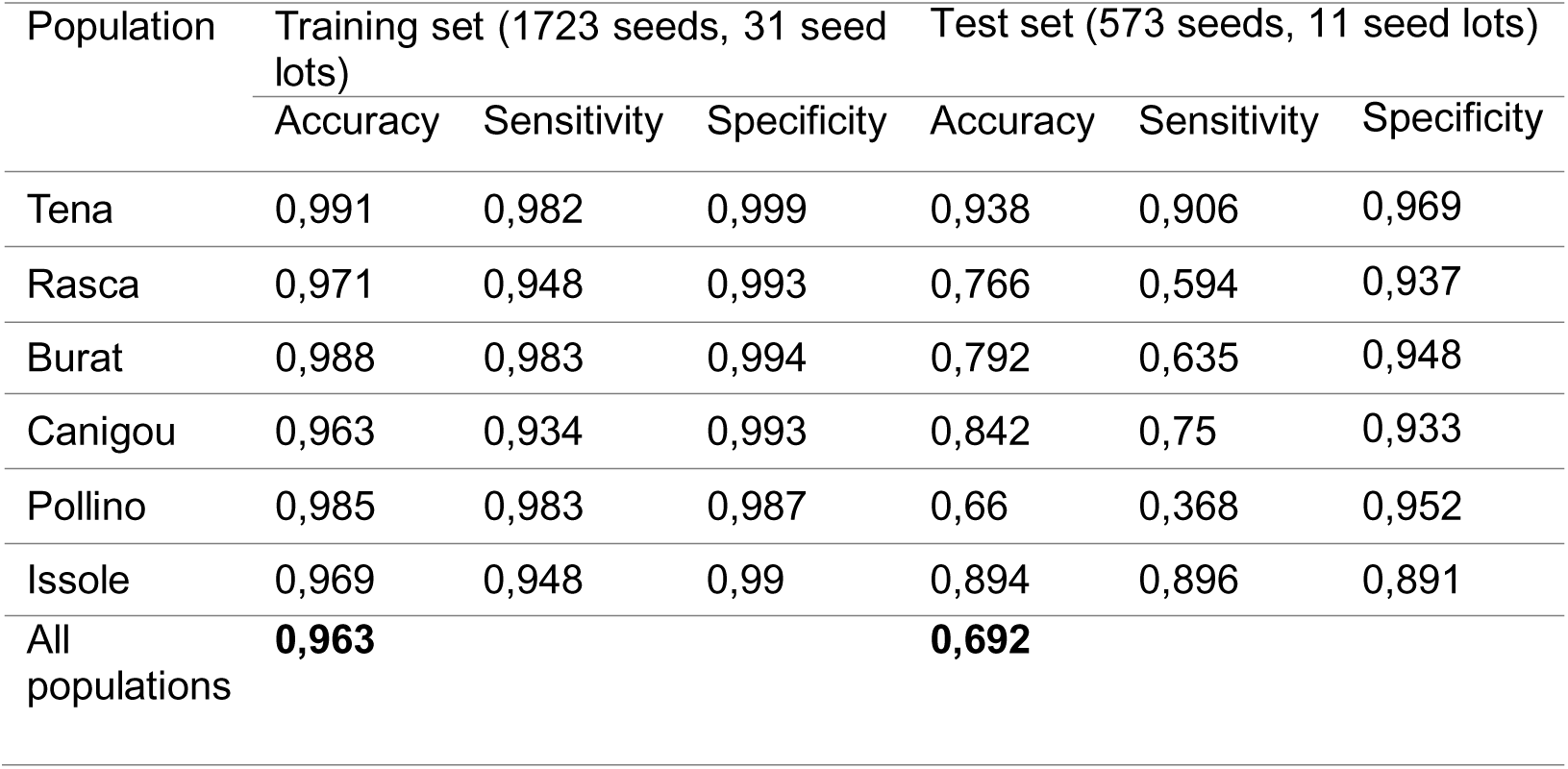
Performances of the PLS-DA model to classify the populations of *A. alba*’s seeds using NIR spectra.

| Population | Training set (1723 seeds, 31 seed lots) |  |  | Test set (573 seeds, 11 seed lots) |  |  |
| --- | --- | --- | --- | --- | --- | --- |
|  | Accuracy | Sensitivity | Specificity | Accuracy | Sensitivity | Specificity |
| Tena | 0,991 | 0,982 | 0,999 | 0,938 | 0,906 | 0,969 |
| Rasca | 0,971 | 0,948 | 0,993 | 0,766 | 0,594 | 0,937 |
| Burat | 0,988 | 0,983 | 0,994 | 0,792 | 0,635 | 0,948 |
| Canigou | 0,963 | 0,934 | 0,993 | 0,842 | 0,75 | 0,933 |
| Pollino | 0,985 | 0,983 | 0,987 | 0,66 | 0,368 | 0,952 |
| Issole | 0,969 | 0,948 | 0,99 | 0,894 | 0,896 | 0,891 |
| All populations | **0,963** |  |  | **0,692** |  |  |

The Canigou and Burat populations from the French Pyrenees tended to be confused with one another (Figure 2): 19% of Burat seeds were attributed to Canigou while 13% of Canigou seeds were attributed to the Burat population. The Pollino seeds that were misclassified were mostly attributed to Issole (34%) and Rasca (18%) populations. The misclassified seeds from Rasca were broadly evenly distributed between the five other populations.

**Figure 2.**
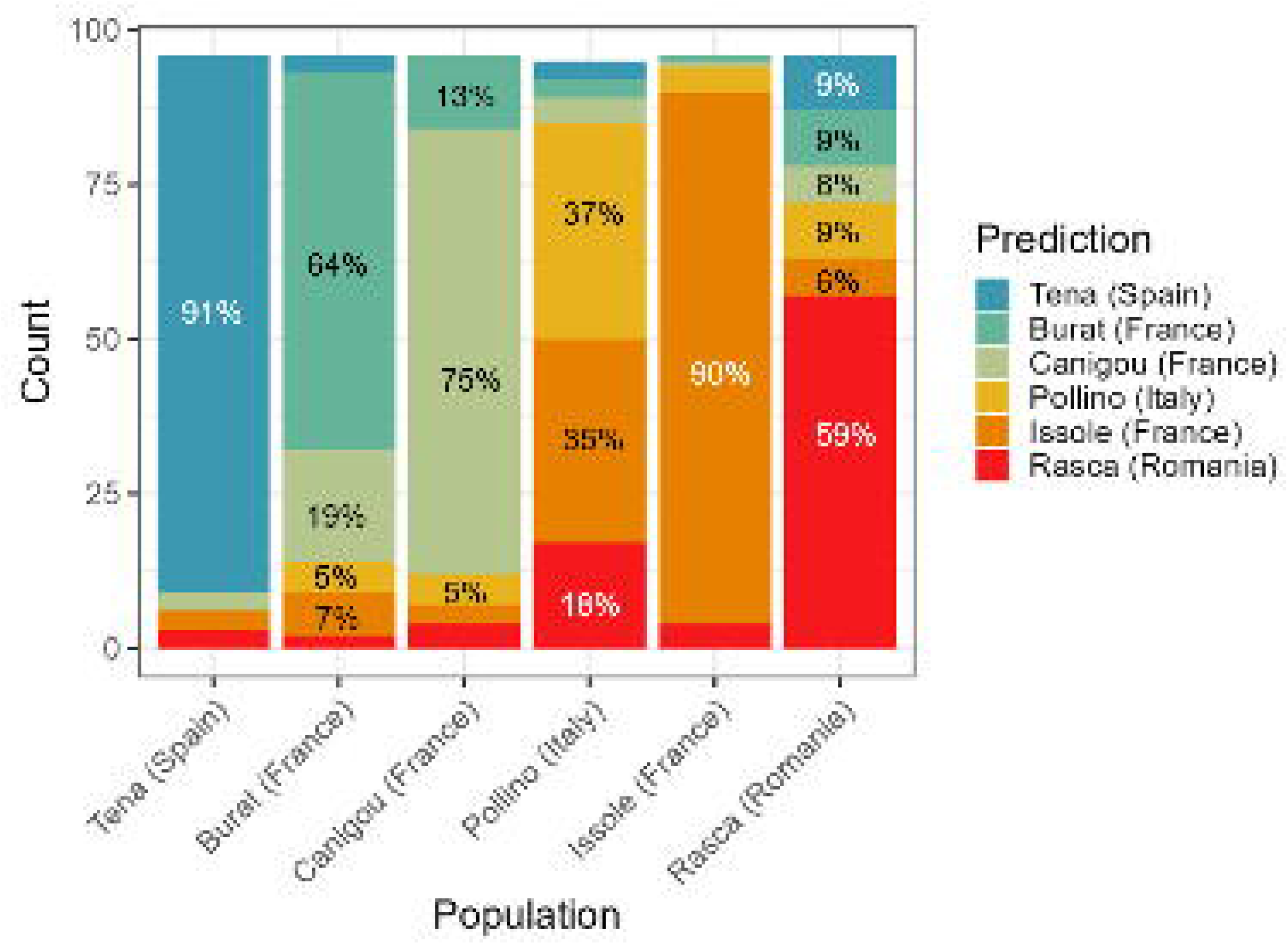
Bar plot of the confusion matrix between the seeds’ actual population and their predicted population by the PLS-DA population model on the external test set (573 seeds, 11 seed lots). Populations are ordered from coldest to warmest maximum spring temperature, with a corresponding blue to red color gradient.

The VIP analysis (Figure 3) revealed which wavelengths were the most important on the separation of the populations. Five main wavebands were the most involved in the seeds’ population prediction: 1404-1419; 1678-1786; 1874-2042; 2094-2208; 2236-2269 (end of spectrum measured). These wavebands encompassed three of the NIR spectra peaks (1712, 1929, 2111 nm).

**Figure 3.**
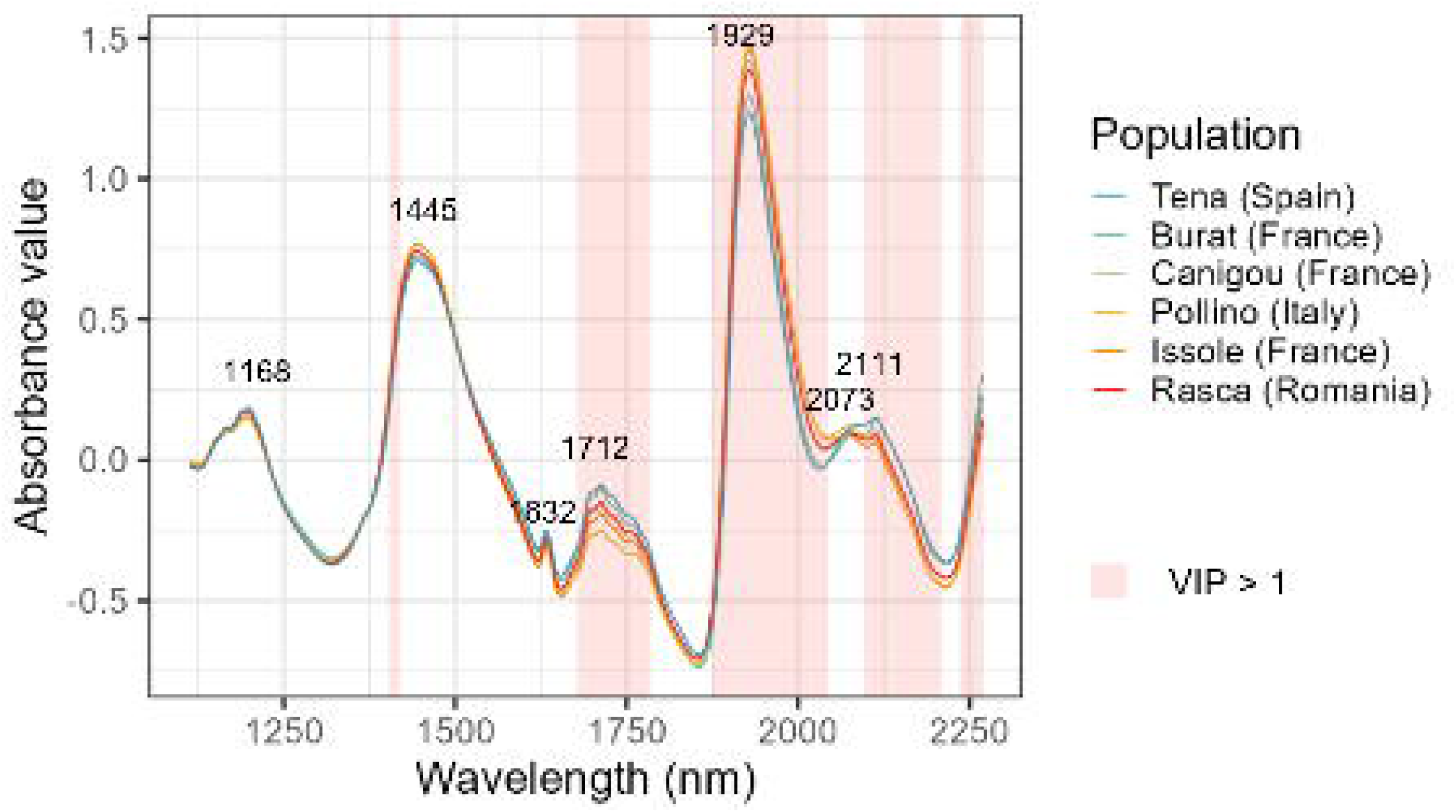
Mean NIR spectra by population with highlights (pink background) of the influential wavelengths (VIP>1) from the PLSDA population model. Populations are ordered from the coldest to the warmest maximum spring temperature, with a corresponding blue to red color gradient.

### 3.2 PLSR to predict the mean germination probability and germination time of the populations

The PLSR germination probability model had 20 latent variables and explained 51% of the variation of germination probability among populations in the test set (Figure 4 a). The PLSR germination time model also had 20 latent variables and explained 65% of the variation of germination time among populations in the test set (Figure 4 b).

**Figure 4.**
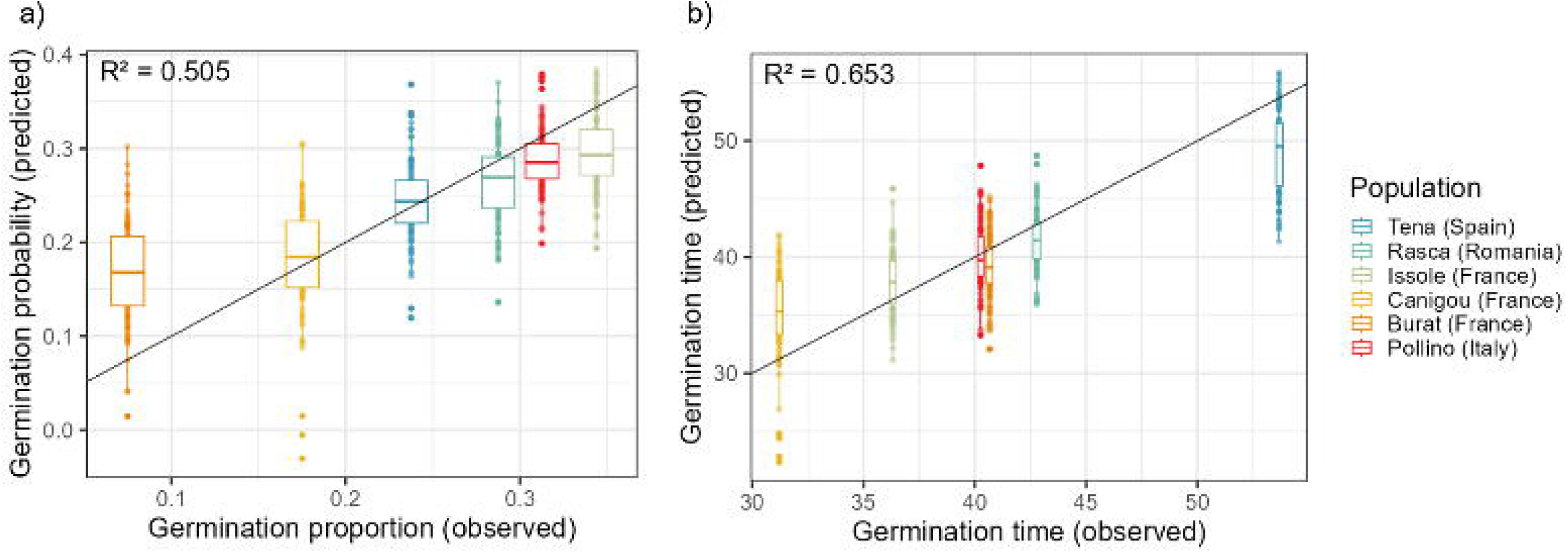
Goodness of fit calculated on the test set for the PLSR germination probability model (a) and the PLSR germination time model (b). The grey line is y = x, i.e., when model prediction is equal to the germination proportion or the mean germination time observed for the population. The population are ordered from coldest to warmest minimum winter temperature, with a corresponding blue to red color gradient.

Median of the predicted values for each population were close to the observed mean, except for the most extreme values: the lowest germination proportion (Burat) and time (Canigou) were overestimated and the highest germination proportion (Issole) and time (Tena) were underestimated.

The VIP analysis highlighted which wavelength were the most important to predict population germination rate and time (Figure 5). The main wavebands that significantly contributed to the construction of the models were respectively: 1403-1424; 1632-1638; 1682-1786; 1846-2033; 2070-2267 nm and 1392-1394; 1398-1411; 1628-1640; 1676-1786; 1871-2033; 2053-2163; 2182-2267.

**Figure 5.**
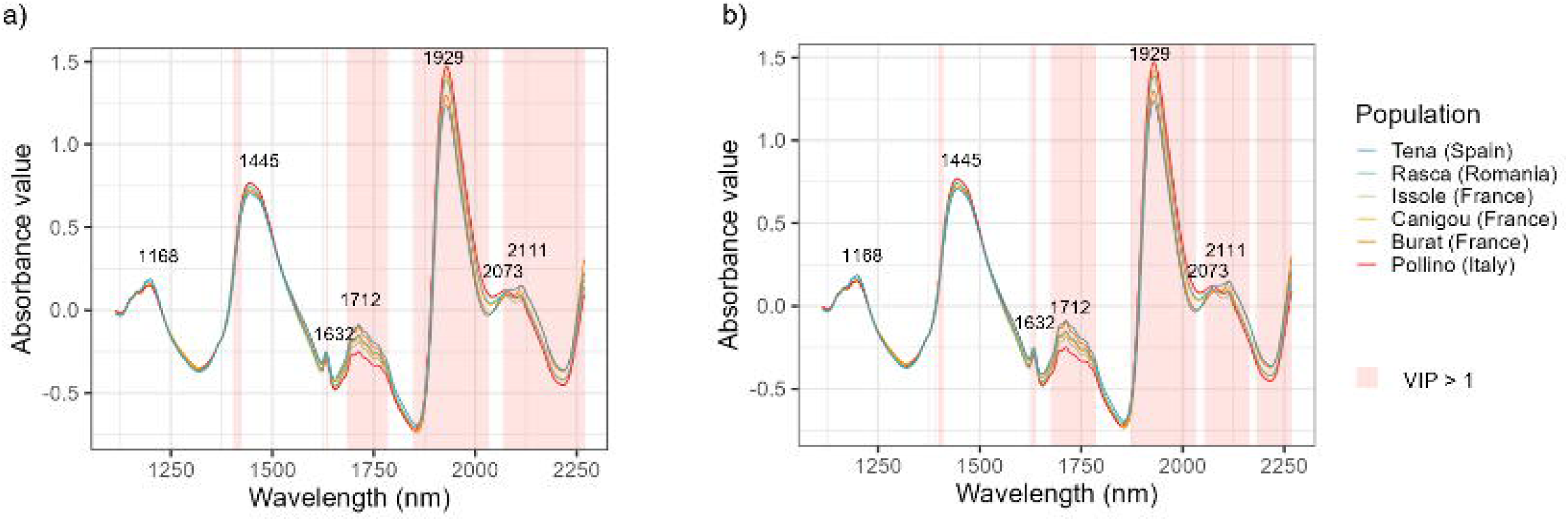
Mean NIR spectra by population with highlights of the influential wavelengths (VIP>1) from the PLSR germination rate model (a) and the PLSR germination time model (b). Populations are ordered from coldest to warmest minimum winter temperature, with a corresponding blue to red color gradient.

These wavebands encompassed the 1712, 1929, 2111 nm peaks, as previously observed for the PLSDA population model, but also included the 1632 and 2073 nm peaks.

### 3.3 GLMM to explain the contribution of population NIR spectra and dormancy release to the germination probability and germination time of seeds

The overall germination rate in the three climatic chambers was 27% (1126 seeds out of 4119 germinated), and the germination time varied from 6 to 132 days (Supplementary Table 4).

Model predictions for germination probability ranged from 0.05 to 0.50.

Dry seed mass (*M*) and NIR population score on the second axis of the PCA (*NIRS*2) had a significant effect probability to germinate. That was not the case for the temperature in the chamber during germination (*CT*), temperature seasonality (*TS*) and stratification treatment (*S*), however their triple interaction on was significant (Supplementary Table S5).

The fixed effects of the model explained 6.1% of the variation in germination probability, and the fixed and random effects together explained 13.1% of the variation. The commonality analysis revealed that *NIRS*2 played the most important role, as it contributed on its own to 43% of the variance explained by the fixed effects of the model (2.6% of the total variance) (Table 3).

**Table 3.** Performances and commonality analyses of germination probability model (binomial general linear mixed model) and germination time model (negative binomial general linear mixed model)

|  | Germination probability |  |  | Germination time |  |  |
| --- | --- | --- | --- | --- | --- | --- |
|  | Marginal | Conditional |  | Marginal | Conditional |  |
| R<sup>2</sup> (trigamma) | 0.0614 | 0.1314 |  | 0.5447 | 0.6025 |  |
| Commonality analysis |  |  |  |  |  |  |
|  | Unique | Common<br>(Sum across<br>all predictor<br>sets) | Total | Unique | Common<br>(Sum across<br>all predictor<br>sets) | Total |
| M | -0.0076 | 0.0157 | 0.0081 | / | / | / |
| NIRS2 | 0.0265 | 0.0358 | 0.0623 | / | / | / |
| TS | 0.0063 | 0.0167 | 0.0230 | / | / | / |
| CT | 0.0001 | 0.0007 | 0.0008 | 0.0000 | 0.3851 | 0.3851 |
| WTmin | / | / | / | 0.0001 | 0.0063 | 0.0064 |
| NIRS1 | / | / | / | 0.0019 | 0.0101 | 0.0120 |
| S | 0.0002 | 0.0000 | 0.0002 | 0.1489 | -0.0029 | 0.1460 |
| CT × WTmin | / | / | / | 0.0000 | 0.0101 | 0.0101 |
| CT × S | -0.0001 | 0.0012 | 0.0011 | 0.0005 | 0.3806 | 0.3811 |
| WTmin × S | / | / | / | 0.0001 | 0.0064 | 0.0065 |
| CT × WTmin × S | / | / | / | 0.0022 | 0.0077 | 0.0099 |
| TS × S | 0.0001 | 0.0230 | 0.0231 | / | / | / |
| TS × CT | 0.0000 | 0.0009 | 0.0009 | / | / | / |
| TS × CT × S | 0.0020 | 0.0014 | 0.0034 | / | / | / |

When adding moisture content (*MC*) to the germination probability model (Supplementary Equation S1), the commonality analysis shows that *NIRS*2 and *MC* have negligible unique effects when both are present, but their shared contribution represented half of the explained variance.

All the variables selected in the germination time model had a significant effect on germination time except for the NIRS population score (*NIRS*1) and minimal Winter Temperature (*WTmin*), however *WTmin*’s interaction with *CT* and the triple interaction with *TS* and *CT* were significant (Supplementary Table S5).

The model predictions ranged from 13 to 98 days to germinate. The fixed effects of the model explained 55% of the variation in germination time, and the fixed and random effects together explained 60% (Table 3).

The commonality analysis revealed that *S* was the predictor with the highest unique contribution (14.9% of the total variance, *i.e.,* 27% of the variance explained by fixed effects) (Table 3). *CT* and the interaction between *CT* and *S* had the highest common effects (0.38 and 0.39), which came almost entirely from the predictor set that included them both, contributing effectively to 71% of the variation explained by the fixed effects of the model. It also confirmed that *NIRS*1’21’s contribution was very small (unique contribution: 0.2%).

According to the model, seeds took less time to germinate if they were stratified and if the temperature during germination was higher (Figure 6). Differences between stratified and non-stratified seeds were larger at cold temperatures. Differences between cold and warm populations were important at 15°C, with seeds from colder population taking longer to germinate, but got weaker when germination temperature increased.

**Figure 6.**
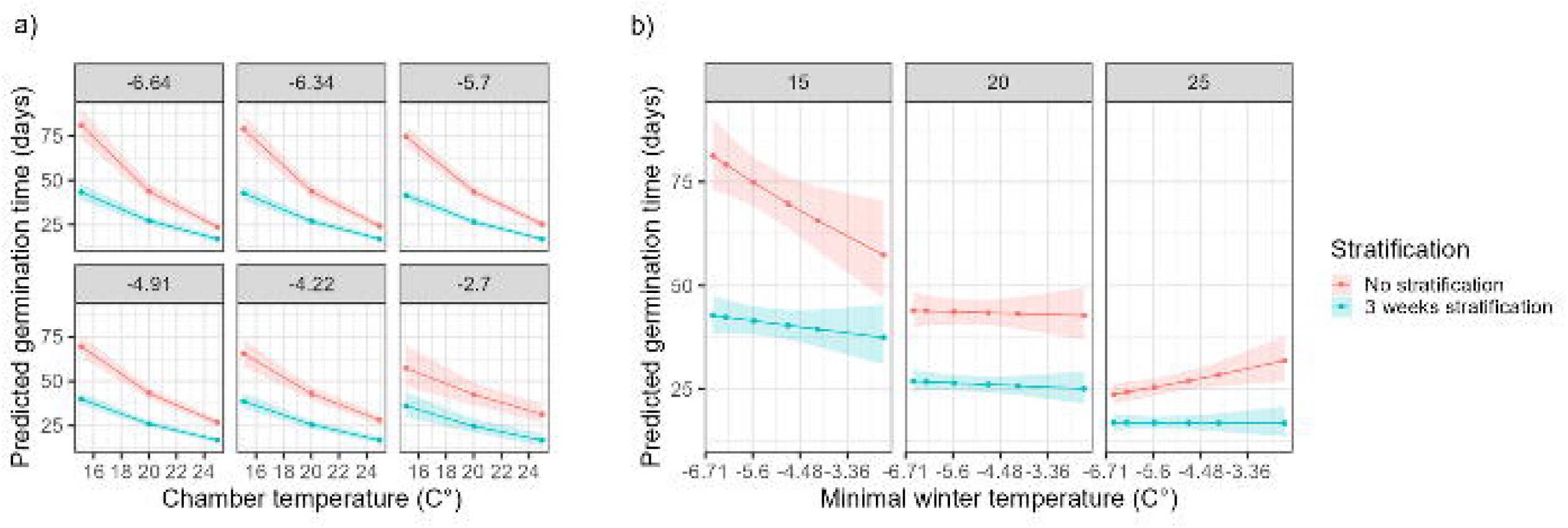
Relationship between germination time, chamber temperature, mean annual temperature at the population origin and stratification from germination time GLMM predictions. Figure a) facets represent the mean annual temperature at the seed population origin and figure b) facets represent the temperature in the chamber during germination. Bootstrapping confidence intervals (95%) are represented by shaded areas.

## 4. Discussion

### 4.1 Population discrimination based on NIRs

The use of NIR spectroscopy to detect seed origin is a promising, non-invasive technique to identify stored bulked seeds. Although it has been widely applied to crop seeds (Choi et al., 2016) and tree nuts (Nisgoski et al., 2023; Vitale et al., 2013), it has been rarely applied in forestry. Farhadi et al. (2017) authenticated the provenance of *Picea abies*, and Tigabu et al. (2005) identified the origin of *Pinus sylvestris* seeds with seeds produced by the same clones among provenances. In these studies, the classification accuracy provided by NIR spectroscopy was close to 90%. Our PSLDA model assigned 69% of the seeds to the correct population in the test set, confirming there are differences in the seed composition between populations of *Abies alba*. Our model had a higher complexity than those used in previous studies, with 35 latent variables. Vitale et al. (2013), for example, had 18 latent variables for as many classes discriminated. Both the differences in classification accuracy and complexity between previous studies and ours could be due to a higher variability of the spectra when measured at a single seed level without considering orientation, and not in bulk or with measurements on both sides of the seed (Vitale et al., 2013).

Our model’s ability to predict seed origin vary between populations and such variation can be explained case by case. The Tena and Issole seeds were well identified by the model (accuracy of 89 and 94%). The Tena and Issole sites showed the lowest and second highest maximum spring temperatures, respectively, and it is likely that this was also the case during the seeds’ development, so their differences in seed composition could be caused by both genetic (adaptation to local climate) and environmental factors during production.

The Canigou and Burat seeds were often confused in the model. This could be due to the geographical and environmental proximity of these sites, which are both located on the north side of the Pyrenees in the reach of gene flow (see **Error! Reference source not found.**). They also share an evolutionary common ancestry (Scotti-Saintagne et al., 2021). If we pooled these two populations together, the model’s performance would go up to 75% (results not shown).

The seeds from Pollino (and to a lesser extent those from the Romanian site of Rasca) were the most misclassified by the model, with particularly low sensitivities. In the studies of Farhadi et al. (2017) and Vitale et al. (2013), poor sensitivity was explained by a higher heterogeneity between seeds, or a higher occurrence of small seeds (whose spectral signals could be weaker). This is not the case for our Rasca and Pollino seeds (Supplementary Figure S1), as they tended to have larger seeds and the variance of the dry weight and moisture content was homogeneous between the groups. Rasca and Pollino were neither the populations with the highest variance in NIR spectra (Supplementary Figure S3). Silver fir populations from southern Italy and the Balkan Peninsula are the most genetically diverse within the species’ distribution. Although they belong to genetic lineages which likely diverged early in Middle Pleistocene, these populations are nowadays genetically similar, possibly due to trans-Adriatic gene flow via-pollen (Liepelt et al., 2002; Piotti et al., 2017). This combination of factors might explain the relatively low predictability shared by these two populations.

It is interesting to note that, according to Liepelt et al., (2009), our six populations all originated from different glacial refugia, with the exception of the Pyrenean populations Tena, Burat and Canigou. Indeed, the genetic structure of silver fir Pyrenean populations allows the identification of three different groups (Scotti-Saintagne et al., 2021). Canigou population belongs to the Eastern Pyrenees group, whereas Burat and Tena likely belong to the Western Pyrenees group. However, our analysis with NIR spectra showed Tena’s population seed spectra very different from the other two Pyrenean populations. These differences between the northern (Bourat and Canigou) and southern (Tena) populations, can be explained by adaptations to the very different conditions occurring at both sides of the Pyrenees. The south side of this mountain range is much sunnier and drier than the north side (Cuadrat et al., 2024)

In summary, environmental conditions and genetic differentiations provide satisfactory explanations to the good performance of the classification of the seeds from the Alps (Issole) and Spain (Tena), as well as the confusion between the seeds from the French Pyrenees (Canigou and Burat). Similarities in the current genetic features of Rasca and Pollino can explain the poor classification seeds coming from these populations.

### 4.2 Prediction of germination properties of population with NIR spectroscopy

Numerous studies have used NIR spectroscopy to discriminate between viable and non-viable seeds in crops (Xia et al., 2019) and trees (Dumont et al., 2015; Tigabu, 2003; Tigabu et al., 2020); however, few have attempted to determine germination probability. Xiao et al. (2023), Çeliktaş and Konuşkan (2020) and Zhang et al. (2023) artificially or naturally aged different seed lots of maize or and rapeseed to simulate a wide range of germination rates. Their models for predicting germination probability showed R² values close to 0.9. In contrast, Al-Amery et al. (2018) used soy-bean seed stored under uniform conditions and durations, yielding R² values between 0.55 and 0.66, which are closer to our R² of 0.51 for the PLSR model on germination probability on same-year seed collection.

To our knowledge, few studies have aimed to predict germination time using NIR spectroscopy. Fan et al. (2020) obtained an accuracy between 90 and 97% to classify germination time of artificially aged wheat kernels, which is more performant than our PLSR model on germination time (R^2^ of 0.65). It is possible that markers of senescence in aged seeds produce stronger NIR signatures, potentially explaining why studies that involved aging had a higher predictive power.

Ultimately, our PLSR models predicted seed germination traits that aligned with the observed mean traits for most populations, confirming that NIR spectroscopy is a suitable tool for identifying seed germination characteristics in forest trees. Calibration with germination data at the mother-tree level, rather than the population level, could further enhance the models’ precision and applicability on a finer scale.

NIR population score, as a proxy for chemical composition, has potential as an explanatory factor to better understand germination. Indeed, when trying to explain germination proportions of individual seeds with GLMM, NIR values of the population was the most important factor and had a significant contribution compared to temperature treatment, climate of origin and seed mass. Temperature treatment and/or climate of origin were yet key predictors in other studies on conifers that did not involve NIR spectroscopy (Hsu et al., 2024; Kueppers et al., 2017; Leinonen et al., 1993). It should be noted that the total variance explained by the GLMM model was rather low (12%), hinting that there could exist other drivers that were not included. Elevation had for instance a significant effect on germination rate in Hsu et al. (2024), although it is generally highly correlated with temperature.

NIR population score contributed very little to explaining germination time with GLMM, the most important factors being stratification and chamber temperature. Considering we had good results with the PLSR on germination time averaged by population using NIR spectroscopy alone, it seems that the specific NIR predictor we used to predict the germination time of each seed in the GLMM was inadequate. The mean score by population on the first axis of the NIR spectroscopy PCA might be too simplistic to explain the subtle mechanisms that drive germination time. A mean score by mother-tree could be a more powerful alternative, but we could not test it due to differences in the mother trees available in the seed lots used for measuring the NIR spectra and germination.

### 4.3 Underlying causes of seed population differentiation and germination based on NIR spectroscopy

NIR spectra are difficult to interpret due to overlapping overtones (Simon, 2020; Workman and Weyer, 2012), thus there is still limited information available on the assignment of bands to specific molecular bonds (Tasumi and Sakamoto, 2015). Nonetheless, several studies have identified important NIR regions in their models associated with key chemical compounds in seeds (Farhadi et al., 2017; Mukasa et al., 2019; Tigabu et al., 2020, 2005; Vitale et al., 2013).

In our study, the VIP analysis in population differentiation, germination probability and germination timing shared peak wavelengths at 1712, 1929 and 2111 nm. The strong absorbance at 1712 nm could be caused by the C-H bound in an aliphatic chain or in an halogenated compound (Workman and Weyer, 2012) whereas the one at 1929 nm could reflect either the absorption by polysaccharides or the presence of water (Workman and Weyer, 2012). Both peak wavelengths have been already similarly interpreted in similar studies of seed provenance differentiation in conifers (Farhadi et al. (2017) and Tigabu et al. (2005)). This would explain why the Tena seeds are very easily identified, as their moisture content is noticeably different from other populations (Supplementary Figure S1). The peak at 2111 nm corresponds to the stretch and bend of a secondary amid according to Chu et al. (2014). Wavebands in the 2100-2500 have been associated with fatty acid (Farhadi et al., 2017 following Hourant et al., 2000; Kim et al., 2007; Ribeiro et al., 2013) and proteins concentrations (Manley, 2014; Workman and Weyer, 2012). These differences found in *Abies alba* NIR spectra among populations suggest variation in polysaccharides, proteins, and fatty acids, which correspond to storage compounds in conifer seeds (Brownfield et al., 2007; Groome et al., 1991). Similar provenance effects on seed composition have been reported in other conifer species (Alvarez et al., 2004; Farhadi et al., 2017). A chemical analysis of the sugar, fatty acid and protein contents of the seeds of the different *Abies alba* populations would be necessary to confirm these interpretations.

Genetic and environmental factors both influence seed composition (Aranda et al., 2012; Fenner, 2010; Piergiovanni et al., 2011) together with maternal effects (developmental characteristics of the mother tree), and the position of the cone in the canopy (Roach and Wulff, 1987). Our sampling design includes enough variation at the mother tree and seed levels to consider that the maternal and cone effects are negligible when looking at differences between populations. Overall, we observed clear population-level differences, especially between Issole and Tena seeds, likely tied to storage compound content. Translocation trials using multi-year seed collections will be needed to disentangle environmental from genetic influences.

In addition to the 1712, 1929 and 2111 nm peaks above described, the VIPs analysis for germination probability and time highlights wavelength peaks at 1632 and 2073 nm. 1632 nm is characteristic of the absorption of a C-H vinyl C-H bound, present in aliphatic hydrocarbons (Workman and Weyer, 2012). 2073 nm corresponds to an absorption band for polypeptides and proteins (Workman, 1996; Workman and Weyer, 2012). These results indicate that determination of germination probability or time is correlated with differences in polysaccharides, moisture, proteins and fatty acids. The importance of moisture is supported by the fact that most of the variance explained by the NIR predictor in the GLMM on germination probability could also be explained by mean moisture content (Supplementary Equation S1). These findings supports other NIR spectroscopy studies on germination (Al-Amery et al., 2018; Çeliktaş and Konuşkan, 2020; Fan et al., 2020; Zhang et al., 2023).

The similarity of influential wavelengths in both population discrimination and germination prediction was expected, given that provenance affects the germination probability of *Abies alba* (Alt, 2022; Boncaldo et al., 2010; Messeri et al., 1963; Morar et al., 2023).

### 4.4 Role of dormancy release in germination at different temperatures of range-wide populations

In temperate biomes, temperature is the most important driver to release physiological dormancy (Penfield and MacGregor, 2017). The extent to which germination will be affected by expected shorter and warmer winters will ultimately determine the chances of tree populations to germinate and regenerate under climate change. Dormancy itself is difficult to measure, as it is linked to the onset of germination, although the duration of stratification provides a reasonably good measurement of the seed dormancy level.

However, we did not detect an effect of stratification on germination probability, as stratification treatment was not retained during GLMM selection. *A. alba* seeds normally present physiological dormancy which, in theory, can lower germination rates if seeds are sown without previous stratification (Wolf, 2003). Our stratification treatment of 3 weeks may not have been enough to provide a complete release of seed dormancy and we acknowledge that longer duration of the stratification period would lead to different results. However, the time of stratification did not affect either the final germination percentage in a germination experiment where Czech silver fir populations have followed different duration in cold stratification, from non-stratification up to 7 weeks (Bezděčková and Řezníčková, 2012). Similar results were found for *Abies cephalonica*, where the cold stratification period did not affect the final germination proportion (Daskalakou et al., 2018). Dormancy release widens hydrothermal thresholds, so it is possible that the temperature gradient tested in our study (and in the other two abovementioned) was too narrow and most populations did not require dormancy release to germinate in these conditions (Fernández-Pascual et al., 2019).

Stratification treatment shortened germination time for every population. The same results where shown by Czech *A. alba* (Bezděčková and Řezníčková, 2012) and Greek *A. cephalonica* populations (Daskalakou et al., 2018), where the time of cold stratification was positively related to the advanced germination. This shows that dormancy release effectively reduced thermal time requirements (Fernández-Pascual et al., 2019). In our experiment, this effect was stronger the colder the chamber was, suggesting that the stratification treatment lowered the minimum temperature allowing germination (Finch-Savage and Leubner-Metzger, 2006). The same was displayed by other species as the invasive grass *Spartina alterniflora* (Cheng et al., 2022), *Aesculus hipocastanum* (Pritchard et al., 1999) and it is a usual response of species with physiological primary dormancy (Baskin and Baskin, 2004). This cold stratification requirement to accelerate germination may preclude winter germination under warming conditions expected by climate change.

Germination time of populations from colder origins was more affected by stratification. This indicates intraspecific variations in dormancy levels, with populations from warmer areas being less dormant than those from colder origin. Such intraspecific variations have been observed in several species such as *Arabidopsis thaliana* (Postma et al., 2016), *Centaurium someadaum* (Fernández-Pascual et al., 2013) or *Aesculus hipocastanum* (Daws et al., 2004), and have been associated to genetic differences between populations as well as to temperature and precipitation variation during seed maturation (lower heat sum or precipitation during maturation inducing deeper primary dormancy). Overall, this result suggests that reduced periods of cold stratification in a climate change context would be more significant for colder populations.

Given our previous ability to discriminate populations based on their NIR spectra, and the correlation between provenance and dormancy levels, it seems feasible to calibrate NIR spectroscopy models to predict dormancy levels among populations. This could be done with NIR spectra measurements and germination experiments involving several stratification treatments. Prediction of seed lots dormancy levels would be especially valuable for afforestation or assisted migration programs.

## Conclusion

Here we showed the value of NIRs to identify seed origins and characterize germination proportions and time in a non-invasive way. This promising approach paves the way to the study of the physiological processes occurring during dormancy, which is of high concern under the new climatic conditions that might produce shorter and warmer winters. Furthermore, it offers a non-destructive tool for assessing seed quality and origin, an invaluable solution for nurseries and the forestry industry, which face the challenge of supplying high-quality, certified seeds of known origin to meet the EU Forest Strategy requirements by 2030.

## Acknowledgements

We thank Cloée Jean for her technical help in setting up the germination experiment. This study was funded by the EU-funded OptForests project (“Harnessing forest genetic resources for increasing options in the face of environmental and societal changes’”; Grant Agreement No 101081774). JP internship was funded by BIOGECO INRAE Unit. MC PhD is funded by the Nouvelle Aquitaine funded project SUBER and the ECODIV division of INRAE.

## Declaration of generative AI and AI-assisted technologies in the writing process

During the preparation of this work the authors occasionally used ChatGPT in order to improve the readability of certain sentences. After using this tool, the authors reviewed and edited the content as needed and take full responsibility for the content of the published article.

## Data statement

All datasets generated during this study will be made available in a public repository upon final acceptance of the manuscript.

