## Supplementary material for "Near-infrared spectroscopy-based models correctly classify *Abies alba* seed origin and predict germination properties": Supplmentary files

Table S1 Number of mother-trees and seeds used in the germination experiment

The number of mother-trees for the Tena population are in brackets because all seeds were pooled together.

| Provenance | Country | No mother-trees | No seeds/modality/mother tree | Total number of seeds |
| --- | --- | --- | --- | --- |
| Issole | France | 8 | 20 | 958 |
| Burat | France | 8 | 10 | 479 |
| Rasca | Rascaania | 8 | 20 | 955 |
| Canigou | France | 8 | 10 | 479 |
| Pollino | Italy | 16 | 3 | 288 |
| Tena | Spain | (20) | 160 (per modality) | 960 |
| Total |  | 48 (+20) |  | 4119 |

Table S2 Number of seeds and mother-trees represented in training and test sets by provenance for the discrimination analysis of the provenances

| Provenance | Training set | | Test set | |
| --- | --- | --- | --- | --- |
|  | Seeds | Mother-trees represented | Seeds | Mother-trees represented |
| Issole | 288 | 6 | 96 | 2 |
| Burat | 288 | 6 | 96 | 2 |
| Canigou | 288 | 6 | 96 | 2 |
| Pollino | 287 * | 6 | 95* | 2 |
| Rasca | 288 | 6 | 96 | 2 |
| Tena | 284 * | / | 96 | / |
| Total | 1723 | 30 (+20 mother-trees from the Tena seed lot) | 575 | 10 (+20 mother-trees from the Spanish seed lot) |

*Four measurements files went missing before analysis and two spectra were excluded because of agarose potentially sliding between the optic fiber and the seeds during measurements.

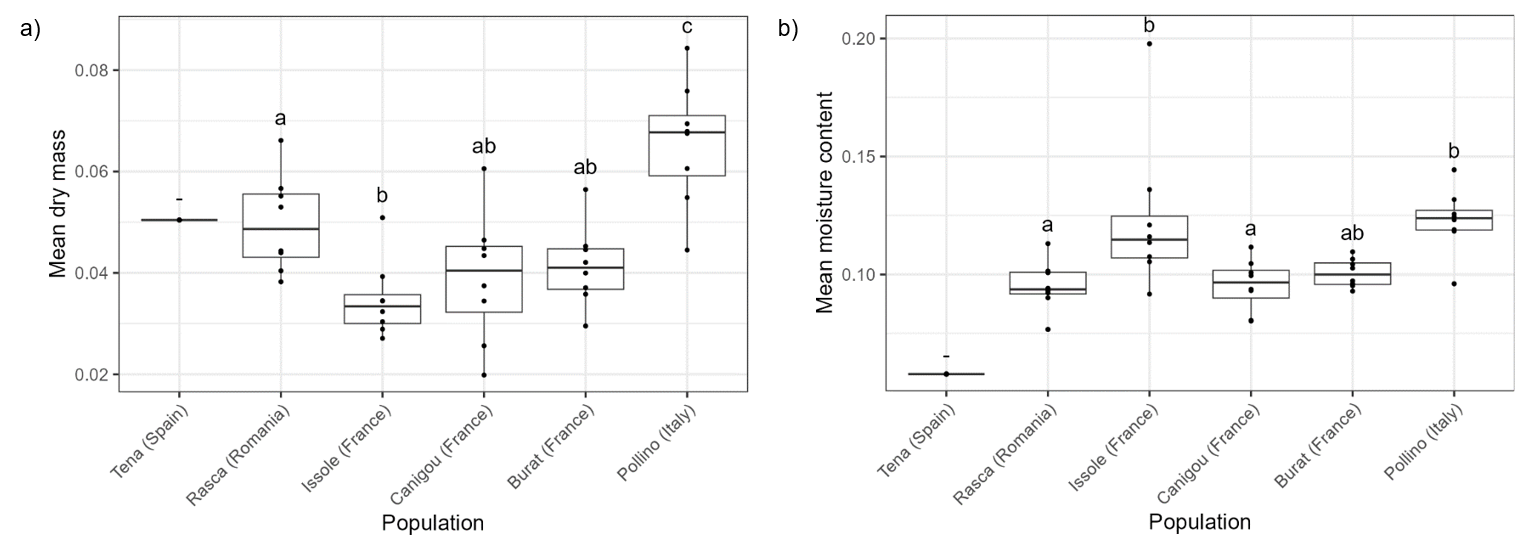

Figure S1 Mean dry mass (a) and mean moisture content (b).

Each point represents a seed lot. There are 8 seed lots in every population except Tena where there is only one. All measures were obtained on twenty seeds. Pairwise comparison labels were obtained with an ANOVA and a Tukey test on all populations except Tena.

### Climate variable selection

To choose the climate variable for each model, a correlation matrix was computed (Table 3) with the germination rate and another with the germination time (averaged at a population level), along with several climate variables describing temperature and precipitation obtained from ClimateDT and averaged over the period 1900-1960 (Marchi et al., 2024): temperature seasonality; De Martonne’s aridity index; mean annual temperature; mean annual precipitation; precipitation sum between April and October; minimal Winter temperature; maximal Spring temperature.

Table S3 Correlation coefficients between germination traits and climate variables

|  | Germination rate | Germination time |
| --- | --- | --- |
| Temperature seasonality | 0.654 | 0.290 |
| De Martonne aridity index | 0.179 | 0.260 |
| Mean annual precipitation | -0.042 | 0.079 |
| Precipitation of driest month | -0.132 | 0.343 |
| Mean annual temperature | 0.122 | -0.163 |
| Precipitation sum between April and October | 0.011 | 0.358 |
| Minimal Winter temperature | -0.397 | -0.524 |
| Maximal Spring temperature | 0.340 | 0.048 |

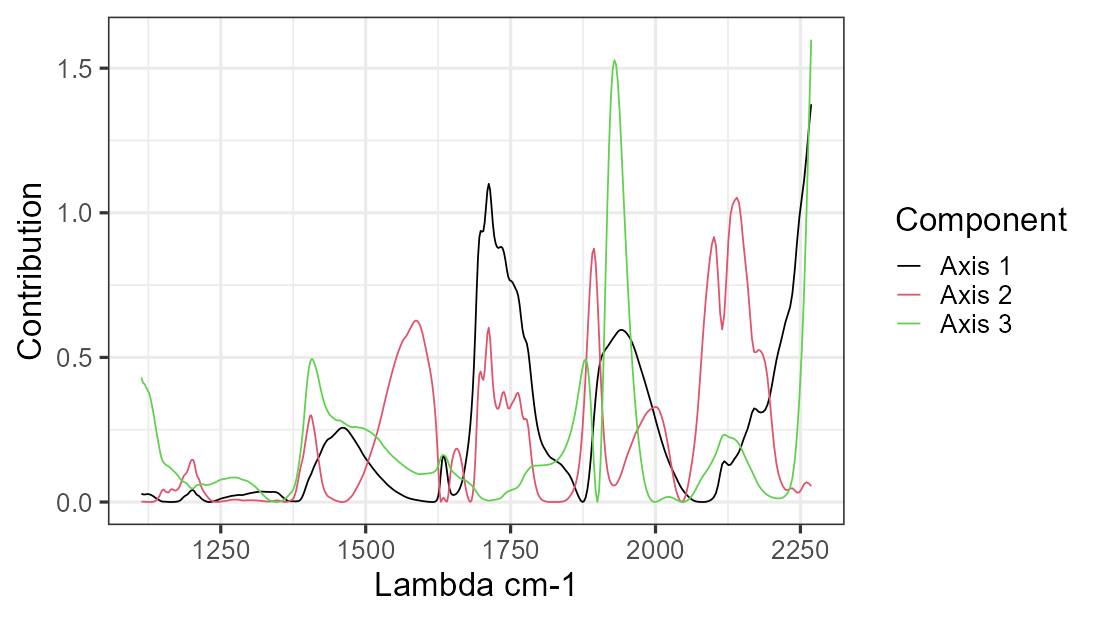

Figure S2 Contribution of each wavelength to the construction of the axes of the PCA of the seeds’ NIR spectra

### Germination rate model with moisture content

The model selection involving moisture content instead of $NIRS2$ lead to the following final model for germination probability:

$$P_{ijk} = \alpha_{0} + \alpha_{1}\cdot M_{jk} + \alpha_{2}\cdot{MC}_{jk} + \alpha_{3}\cdot{TS}_{k} + \alpha_{4}\cdot{CT}_{ijk} +\alpha_{4}\cdot S_{ijk} +$$

$${\begin{aligned} \alpha\end{aligned}}_{5}\cdot TSk \times CTijk + \alpha6\cdot TSk \times Sijk+ \alpha7\cdot CTijk \times Sijk+$$

$$\alpha_{8}\cdot TSk \times CTijk \times Sijk + \beta+ \varepsilon$$

(S1)

With $P_{ijk}$ the probability the *i*th seed of the *j*th mother-tree of the *k*th provenance will germinate; $M_{jk}$ the mean seed mass of the *j*th mother tree of the *k*th population; ${MC}_{jk}$ the mean moisture content of the *j*th mother tree of the *k*th population; ${TS}_{k}$ the temperature seasonality of the *k*th population; ${CT}_{ijk}$ the day temperature of the climatic chamber in which the *i*th seed of the *j*th mother-tree of the *k*th provenance was cultivated; $S_{ijk}$ the stratification treatment (binary) of *i*th seed of the *j*th mother-tree of the *k*th provenance; $\beta$ the random effects and $\varepsilon$ the residuals.

$M$, $MC$, and the triple interaction between $TS$, $CT$ and $S$ had a significant effect on the probability to germinate. The fixed effects of the model explained 5.5% of the total variance, and 12% with the random effects. According to the contribution analysis, moisture content was the highest contributor (37% of the variance explained by fixed effects). When adding $NIRS2$ in the model and running a second contribution analysis, $MC$ and $NIRS2$ ‘s unique contributions were respectively -0.001 and -0.0072, and the shared contribution of the predictor set with them both was 0.028 (51% of the variance explained by fixed effects).

Replacing $NIRS2$ with $MC$ in the germination time model selection does not change the final model.

Table S4 Germination characteristics, mean moisture content and dry seed mass for each seed provenance and modality used in the germination assay

| Population | | Stratification | Chamber temperature | Number of seeds | Germination proportion | Minimum germination time | Maximum germination time | Mean germination time | Mean moisture content | Mean dry seed mass |
| --- | --- | --- | --- | --- | --- | --- | --- | --- | --- | --- |
| ISSOLE | 0 | | 15 | 160 | 0.31875 | 21 | 21 | 75.72549 | 0.0712 | 0.046059 |
| ISSOLE | 0 | | 20 | 160 | 0.34375 | 10 | 10 | 36.30909 | 0.0712 | 0.046059 |
| ISSOLE | 0 | | 25 | 160 | 0.29375 | 10 | 10 | 26.06383 | 0.0712 | 0.046059 |
| ISSOLE | 1 | | 15 | 160 | 0.3125 | 19 | 19 | 53.64 | 0.0712 | 0.046059 |
| ISSOLE | 1 | | 20 | 159 | 0.220126 | 8 | 8 | 28 | 0.0712 | 0.046054 |
| ISSOLE | 1 | | 25 | 159 | 0.301887 | 6 | 6 | 17.64583 | 0.0712 | 0.04603 |
| BURAT | 0 | | 15 | 80 | 0.175 | 39 | 39 | 76.14286 | 0.0856 | 0.051006 |
| BURAT | 0 | | 20 | 80 | 0.075 | 19 | 19 | 40.66667 | 0.0856 | 0.051006 |
| BURAT | 0 | | 25 | 80 | 0.2 | 10 | 10 | 31.3125 | 0.0856 | 0.051006 |
| BURAT | 1 | | 15 | 80 | 0.125 | 19 | 19 | 50.7 | 0.0856 | 0.051006 |
| BURAT | 1 | | 20 | 80 | 0.125 | 19 | 19 | 35 | 0.0856 | 0.051006 |
| BURAT | 1 | | 25 | 79 | 0.101266 | 8 | 8 | 15.75 | 0.0856 | 0.051 |
| CANIGOU | 0 | | 15 | 80 | 0.3125 | 24 | 24 | 64.04 | 0.0794 | 0.038826 |
| CANIGOU | 0 | | 20 | 80 | 0.175 | 17 | 17 | 31.21429 | 0.0794 | 0.038826 |
| CANIGOU | 0 | | 25 | 80 | 0.15 | 14 | 14 | 26.33333 | 0.0794 | 0.038826 |
| CANIGOU | 1 | | 15 | 80 | 0.2125 | 19 | 19 | 39.35294 | 0.0794 | 0.038826 |
| CANIGOU | 1 | | 20 | 80 | 0.2375 | 12 | 12 | 34.36842 | 0.0794 | 0.038826 |
| CANIGOU | 1 | | 25 | 79 | 0.139241 | 6 | 6 | 16.72727 | 0.0794 | 0.038803 |
| POLLINO | 0 | | 15 | 48 | 0.083333 | 46 | 46 | 80.75 | 0.0758 | 0.075163 |
| POLLINO | 0 | | 20 | 48 | 0.3125 | 11 | 11 | 40.26667 | 0.0758 | 0.075163 |
| POLLINO | 0 | | 25 | 48 | 0.479167 | 12 | 12 | 32.21739 | 0.0758 | 0.075163 |
| POLLINO | 1 | | 15 | 48 | 0.333333 | 19 | 19 | 32.125 | 0.0758 | 0.075163 |
| POLLINO | 1 | | 20 | 48 | 0.270833 | 12 | 12 | 20.46154 | 0.0758 | 0.075163 |
| POLLINO | 1 | | 25 | 48 | 0.25 | 8 | 8 | 16.83333 | 0.0758 | 0.075163 |
| RASCA | 0 | | 15 | 160 | 0.35 | 24 | 24 | 91.03571 | 0.0698 | 0.054527 |
| RASCA | 0 | | 20 | 160 | 0.2875 | 12 | 12 | 42.78261 | 0.0698 | 0.054527 |
| RASCA | 0 | | 25 | 160 | 0.375 | 10 | 10 | 25.71667 | 0.0698 | 0.054527 |
| RASCA | 1 | | 15 | 157 | 0.388535 | 22 | 22 | 49.56667 | 0.0698 | 0.054457 |
| RASCA | 1 | | 20 | 158 | 0.360759 | 12 | 12 | 25.80702 | 0.0698 | 0.054663 |
| RASCA | 1 | | 25 | 160 | 0.19375 | 6 | 6 | 18.83871 | 0.0698 | 0.054527 |
| TENA | 0 | | 15 | 160 | 0.33125 | 46 | 46 | 102.4717 | 0.0739 | 0.065139 |
| TENA | 0 | | 20 | 160 | 0.2375 | 17 | 17 | 53.66667 | 0.0739 | 0.065139 |
| TENA | 0 | | 25 | 160 | 0.25 | 10 | 10 | 24.575 | 0.0739 | 0.065139 |
| TENA | 1 | | 15 | 160 | 0.21875 | 15 | 15 | 56.05714 | 0.0739 | 0.065139 |
| TENA | 1 | | 20 | 160 | 0.425 | 12 | 12 | 23.94118 | 0.0739 | 0.065139 |
| TENA | 1 | | 25 | 160 | 0.3125 | 8 | 8 | 19.26 | 0.0739 | 0.065139 |

Table S5 Fixed and random effects of germination probability response (binomial general linear mixed model) and germination time response (negative binomial general linear mixed model). Significance codes: 0 ‘***’ 0.001 ‘**’ 0.01 ‘*’ 0.05 ‘.’ 0.1 ‘ ’ 1

|  | Germination probability | | | Germination time | | |
| --- | --- | --- | --- | --- | --- | --- |
|  | Estimate | P-value | Significance | Estimate | P-value | Significance |
| Random effects (mother tree) | | | | | | |
| Variance | 0.265 | / | / | 0.02253 | / | / |
| Standard deviation | 0.5148 | / | / | 0.1501 | / | / |
| Fixed effects | | | |  |  |  |
| Intercept | -1.21732 | < 2e-16 | *** | 3.780076 | < 2e-16 | *** |
| M | 0.16876 | 0.02143 | * | / | / | / |
| NIRS2 | 0.38291 | 0.00240 | ** | / | / | / |
| TS | 0.19755 \| \| . | 0.05058 | . | / | / | / |
| CT | -0.02005 | 0.69177 |  | -0.438910 | < 2e-16 | *** |
| S | -0.05479 | 0.44687 |  | -0.492828 | < 2e-16 | *** |
| WTmin | / | / | / | -0.004047 | 0.88372 |  |
| NIRS1 | / | / | / | 0.048610 | 0.21056 |  |
| CT × S | -0.05910 | 0.41222 |  | 0.075914 | 0.00192 | ** |
| CT × WTmin | / | / | / | 0.072132 | 1.88e-05 | *** |
| S × WTmin | / | / | / | -0.017138 | 0.49653 |  |
| CT × WTmin × S | / | / | / | -0.054533 | 0.03090 | * |
| TS × CT | 0.04036 \| \| | 0.41494 |  | / | / | / |
| TS × S | -0.04484 | 0.52818 |  | / | / | / |
| TS × S × CT | -0.19286 | 0.00679 | ** | / | / | / |

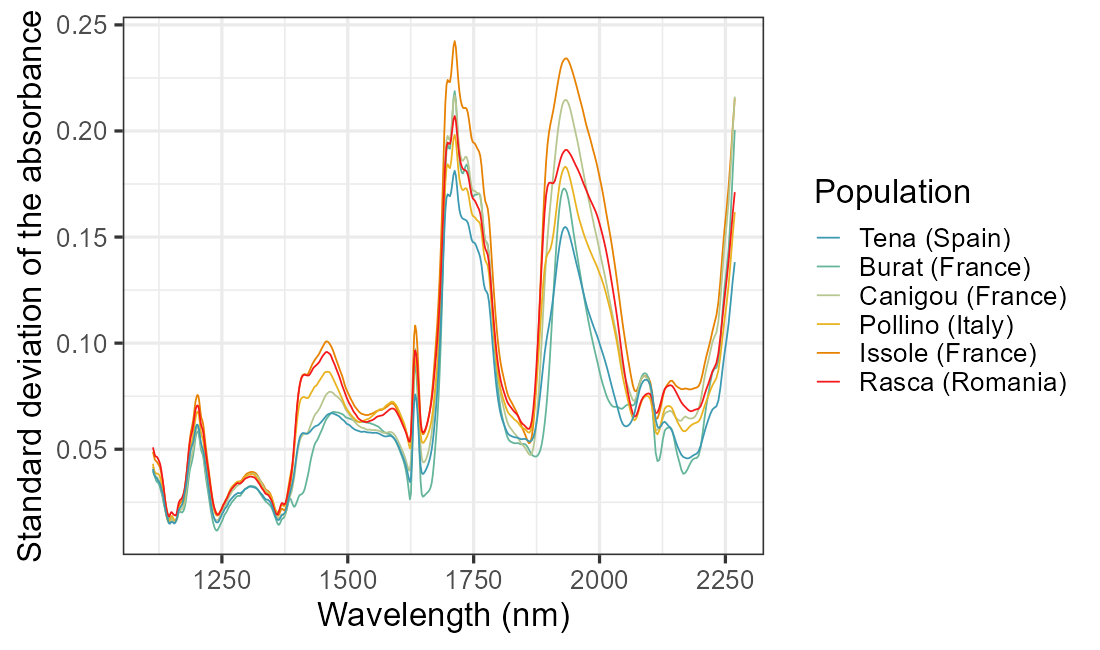

Figure S3 Standard deviation of the NIR spectra for each population

Marchi. M.. Bucci. G.. Iovieno. P.. Ray. D.. 2024. ClimateDT: A Global Scale-Free Dynamic Downscaling Portal for Historic and Future Climate Data. Environments 11. 82. https://doi.org/10.3390/environments11040082
